# Recombinant *Fasciola hepatica* fatty acid binding protein (Fh15) as a novel anti-inflammatory biotherapeutic in an acute gram-negative non-human primate sepsis model

**DOI:** 10.1101/2021.09.21.461321

**Authors:** Jose J. Rosado-Franco, Albersy Armina-Rodriguez, Nicole Marzan-Rivera, Armando G. Burgos, Natalie Spiliopoulos, Stephanie M. Dorta-Estremera, Loyda B. Mendez, A.M. Espino

**Author notes:** **Corresponding Author** Ana M. Espino, Department of Microbiology and Medical Zoology; University of Puerto Rico-Medical Sciences Campus, PO BOX 365067, San Juan, PR 00936-5067, /1318.

## Abstract

Due to their phylogenetic proximity to human, non-human primates (NHP) are considered an adequate choice for basic and pre-clinical model of sepsis. Gram-negative bacteria are the primary causative of sepsis. During infection bacteria continuously release the potent toxin lipopolysaccharide (LPS) into the bloodstream, which triggers an uncontrolled systemic inflammatory response leading to death. Our previous research has demonstrated *in vitro* and *in vivo* using a mouse model of septic shock that Fh15, a recombinant variant of the *Fasciola hepatica* fatty acid binding protein, acts as an antagonist of TLR4 suppressing the LPS-induced pro-inflammatory cytokine storm. The present study aimed to demonstrate that Fh15 suppress the cytokine storm and other inflammatory markers during the early phase of an endotoxemia induced in rhesus macaques by i.v. infusion with lethal doses of live *E. coli*. Fh15 was administrated as isotonic infusion 30 min prior to the bacterial infusion. Among the novel findings reported in this communication, **I)** Fh15 significantly prevented bacteremia, suppressed LPS levels in plasma and the production of C-reactive protein and procalcitonin, which are key signature of inflammation and bacterial infection, respectively, **II)** notably reduced the production of pro-inflammatory cytokines, and **III)** increased innate immune cell populations in blood, which suggest a role in promoting a prolonged steady state in rhesus macaques even in the presence of inflammatory stimuli. This is the first report demonstrating that a *F. hepatica*-derived molecule possesses potential as anti-inflammatory drug against endotoxemia in an NHP-model.

**Tweet:** This is the first communication demonstrating that a *F. hepatica*-derived molecule possesses potential as anti-inflammatory drug against endotoxemia in an NHP-model.

**Importance:** Sepsis caused by Gram-negative bacteria affect 1.7 million adults annually in the United States and is one of the most important causes of death at intensive care units. Although the effective use of antibiotics has resulted in improved prognosis of sepsis, the pathological and deathly effects has been attributed to the persistent inflammatory cascade. There is a present need to develop anti-inflammatory agents that can suppress or neutralize the inflammatory responses and prevent the lethal consequences of sepsis. We demonstrated herein that a small molecule of 14.5kDa can suppress the bacteremia, endotoxemia and many other inflammatory markers in a rhesus macaque model. These results reinforce the notion that Fh15 constitute an excellent candidate for drug development against sepsis.

## Introduction

As part of their immunomodulatory mechanisms, helminths establish a regulatory anti-inflammatory immune response in their mammalian host with a prominent T helper-2/ T regulatory (Th2/Treg) immune profile (1, 2), which is thought to be mutually beneficial for host and parasite, because it protects the host from severe consequences of inflammatory responses while preventing the elimination of worms (3). Thus, a large number of human and animal studies have demonstrated that helminth infections could be used to ameliorate or prevent inflammatory diseases (4-8). In fact, severe sepsis is significantly less frequent in persons carrying chronic helminth infections than it is in non-parasitized persons (1). These studies have helped incorporate helminth infections into the expanded ‘Hygiene hypothesis’ (9). *Fasciola hepatica*, one of the most prevalent parasitic Platyhelminths, is not an exception. However, because of the pathogenicity and negative impact that *F. hepatica* exerts on the health of animals and humans, the infection with this parasite cannot be used to treat inflammatory diseases in humans. Additionally, the immune regulation associated with *F. hepatica* infection lacks specificity and results in a compromised immune system unable to respond effectively to bystander infections (10, 11). We consider it more judicious to identify and purify defined immune-modulatory molecules produced by the parasite, which have the potential for drug development and to characterize their precise mechanism of action. Owing to its extraordinary capabilities for the host’s immune-modulation, *F. hepatica* constitutes an enormous ‘pharmacopeia’. As soon as the parasite invades the gut wall, it initiates a complex interaction with various host immune cells (e.g. macrophages or dendritic cells). The parasite secretes a myriad of immunomodulatory molecules termed excretory-secretory products (ESPs) that direct the host’s immune response toward a non-protective Th2/Treg environment with suppressed Th1 immunity, which allows the parasite to persist in the host for a long period of time (12, 13). Some of these molecules belong to the fatty acid binding protein family (FABP).

*F. hepatica* FABPs are known antioxidant molecules (14) that have been extensively used as vaccine candidates against fascioliasis or schistosomiasis (15-17). In a previous study we demonstrated that a single therapeutic dose of recombinant *F. hepatica* FABP (Fh15; 50μg) given to mice 1h after exposure to lethal LPS challenge significantly suppressed pathological sequelae by concurrently modulating the dynamic of macrophages in the peritoneal cavity and the activation status of spleen macrophages in a mouse model of septic shock (18).

Since non-human primates (NHP) share physiological and anatomical features similar to humans and have shown to respond similar to humans when exposed to live Gram-negative bacteria (19), they represent a more relevant pre-clinical model than rodents-model to study the inflammatory responses during the acute phase of sepsis. With these advantages in mind, in a previous study we developed a rhesus macaque model of septic shock induced by intravenous (i.v.) administration of live *Escherichia coli* to identify inflammation-associated markers during the early phase of sepsis in rhesus macaques (20). As a result of that study, we were able to determine that bacteremia was present in all animals from 30 minutes to 3 hours following *E. coli* infusion whereas endotoxin, C-reactive protein (CRP) and Procalcitonin (PCT) were detected during the full-time course suggesting an ongoing inflammatory process caused by an active bacterial infection. Similarly, TNF-α was detected at 2h whereas IL-6, IL-12, and IFN-γ were detected after 4h of *E. coli* infusion (20). The present study aims to assess whether these inflammatory markers can be suppressed when Fh15 is administered i.v. as isotonic infusion 30 min before a challenge with live *E. coli*-infusion. The present study is the first to demonstrate that Fh15 has the capacity to suppress bacteremia, endotoxemia, CRP, PCT and pro-inflammatory cytokines and chemokines during the early phase of an *E. coli*-induced acute sepsis in a rhesus macaque model, in addition to modulating the population of innate immune cells in blood.

## Materials and Methods

### Ethics statement

This study was performed in accordance with the Guide for the Care and Use of Laboratory Animals (National Research Council (US), 2011) and was approved by the Institutional Animal Care and Use Committee of the University of Puerto Rico-Medical Sciences Campus (Protocol No. 7870116).

### Animals

Nine adult male (6-7 years-old) rhesus macaques (*Macaca mulatta*), weighing 6.4-9.66kg (8.1±1.08) were facilitated by the Caribbean Primate Research Center at the University of Puerto Rico-Medical Sciences Campus. Prior to inclusion in the experiment, the monkeys received a physical examination and were tested for hematological, serological, and microbiological abnormalities. Only the animals confirmed healthy by a veterinarian were included.

### High-throughput protein expression and purification of Fh15

cDNA encoding full length fatty acid binding protein from *F. hepatica* (Fh15) (GenBank: M95291.1) (21) was synthesized and cloned into the pT7M vector and expressed as fusion protein with His-Tag (6xHis) at amino terminal in a novel bacterial expression system, using *Bacillus subtilis* (Genscript USA). This model offers more advantages than our previously optimized Fh15-expression system in *E. coli* (22). *B. subtilis* is a non-pathogenic gram-positive bacterium that does not produce LPS. Furthermore, it is not codon biased, grows faster, and has higher secretory capacity. *Bacillus* strain 7024E was transformed with recombinant plasmid. A single colony was inoculated into TB medium; culture was incubated at 37°C and when the OD_600_ reached about 1.2, protein overexpression was induced with IPTG at 37°C for 4h. Cells were harvested by centrifugation and cell pellets were resuspended with lysis buffer followed by sonication. The precipitate after centrifugation was dissolved using denaturing agent. Fh15 was purified from inclusion bodies by one-step purification using Ni-column. Fh15 was stabilized in PBS containing 10% Glycerol, 0.5M NaCl, and pH 7.4 and sterilized via a 0.22μm filter. Western blot using a mouse anti-Histidine tag monoclonal Antibody (Genscript Cat. No. A00186) was used to confirm purity of the purified protein (**Figure-1S**). Purified Fh15 had endotoxin levels lower than 0.2 EU/mg measured by LAL Endotoxin Assay kit (Bioendo, Cat. No. KC64T). Protein concentration of Fh15 (3.86 mg/ml) was determined by Bradford method with BSA as a standard (ThermoFisher Cat. No. 23236).

### Anti-Fh15 polyclonal antibody and conjugation

Polyclonal antibody against Fh15 was produced in New Zealand White rabbits by subcutaneous injections of 200μg of protein mixed with an equal amount of complete Freund’s adjuvant in the first injection, and incomplete Freund’s adjuvant in the boost injections as previously described (23). Anti-serum had antibody titers of ∼1:100,000 when was titrated by indirect ELISA against the Fh15. Anti-Fh15 polyclonal IgG was purified by affinity chromatography using 5/5/HiTrap Protein-A HP (GE Healthcare, Piscataway, NJ). The resulting IgG was conjugated with horseradish peroxidase (HRP, Abcam, UK, ab102890) and used to develop a Sandwich ELISA to detect the presence of circulating Fh15 in the blood of rhesus macaques later used in this study.

### Escherichia coli culture

The *Escherichia coli* 086a: K61 serotype used to induce sepsis was purchased from American Type Culture Collection (ATCC, 33985). Two days prior to each experiment a fresh glycerol stock was plated on a Luria Broth agar and cultured for 20 hours at 37°C. The next day a single colony was cultured in 300ml of Luria Broth for 16-18 hours at 37°C, until it reached a concentration of 10^10^ CFU/ml. Afterwards, to remove free LPS the culture was harvested and washed twice with endotoxin-free PBS (0.01M phosphate buffered saline pH 7.4). The pellet was resuspended in 50ml of endotoxin-free PBS and administrated to animals for 2h.

### Animal preparation, and intravenous infusion administration

Animals were fasted overnight and sedated the next day with ketamine hydrochloride (100 mg/ml) intramuscularly (10 mg/kg) (Akorn-Lake Forrest, IL). Afterwards, they were transported to the surgical suite and anesthetized with isoflurane gas (Akorn Inc-Lake Forest, IL) via facemask. Then animals were intubated with 3.0 -3.5 mm endotracheal tube for maintenance anesthesia and were kept intubated for the entire 8-hour period allowing them breath on their own. A 20-gauge intravenous catheter (Nipro-Osaka, Japan) was inserted in the lateral saphenous to inject the infusions prepared in isotonic saline containing 2.5% dextrose at a rate of 3.3 ml/kg/h to compensate fluid loss. To avoid discomfort, the animals remained under anesthesia during the entire experimental window of 8-hours. Animals were connected to a BIONET monitor that measured physiological parameters such as body temperature (BT), heart rate (HR), respiratory rate (RR), and mean arterial blood pressure (MAP) every 10 minutes. The animals remained under constant monitoring by veterinary staff during the complete time-course of the experiment.

The bacterial infusion consisted of 50-ml isotonic saline solution containing 10^10^ CFU/kg body weight (wt.) of live *E. coli*, which was applied at a constant rate infusion (CRI) of 0.42ml/min for 2h. This dose was determined to be a lethal dose for these subjects as reported elsewhere (24). The Fh15-infusion consisted of 5ml isotonic saline containing 12mg Fh15 applied at a constant rate infusion of 0.25ml/min for 20 minutes. Because there are no previous studies with rhesus monkeys that have been injected i.v. with proteins, the Fh15 dose administered in this study was arbitrarily selected. The Fh15 administered was equivalent to 1.24 to 1.87mg/kg of body weight (average 1.48mg/kg body wt.).

### Experimental groups

Naïve rhesus monkeys were randomly allotted into four experimental groups (n=3/group) designated as *E. coli*, Fh15, Fh15-*E. coli* and Fh15-Fh15-*E. coli*. The group designated, as *E. coli* was a positive control group for sepsis that only received the bacterial isotonic infusion. Following *E. coli*-infusion, animals were monitored for 8h and then euthanized. The group designated as Fh15 only received an isotonic infusion of 5ml containing 12mg Fh15 for 20min and after monitored for 8h they were allowed to recover from anesthesia and returned to their original cages for 3 months. The group named Fh15-*E. coli* received the isotonic infusion of Fh15 for 20min followed immediately by an *E. coli* infusion and were euthanized at 8h of experimentation. The group named as Fh15-Fh15-*E. coli* comprised the same animals that three months before had received the Fh15-infusion. After that lag-time these animals received a second Fh15-infusion (12mg) followed immediately by the lethal *E. coli*-infusion and were euthanized at 8h of experimentation as described above **(Figure-2S)**.

Blood samples of 5ml were taken from the femoral vein of all animals using a 20-gauge Vacutainer needle into heparinized tubes at 0, 30 min, 2h, 4h, 6h and 8h of experimentation. Blood samples were centrifuged at 10,000 rpm × 10 minutes and the plasma was collected and stored at −20°C in aliquots of 500μl until further use. Prior to centrifugation, aliquots of 500μl and 150μl from each blood sample were allotted to determine bacteremia levels and quantifying immune cell populations by flow cytometry, respectively. After the completion of the experimental window, animals were euthanized with pentobarbitol solution 390 mg/ml (Med-Pharmex-Pomona, CA) according the *AVMA Guidelines for the Euthanasia of Animals* (2013). Post-mortem examination of all animals was conducted immediately after animals were euthanized. Gross necropsy was performed and examined for any abnormalities.

### Bacteremia and endotoxin levels assessment

To assess the number of viable bacteria, 500μl of whole blood was diluted 1:1 with sterile 1xPBS, spread in a Luria Broth (LB) agar plate and culture for 20h at 37°C. Colonies we counted and adjusted by the dilution factor. The presence of circulating levels of LPS was determined using Pierce™ Endotoxin Quant kit following manufacturer’s instruction (Thermo Research Scientific, US, A39553).

### C-Reactive Protein (CRP) and Procalcitonin (PCT) concentration determination

Plasma samples from each time point were tested for the quantification of CRP and PCT, which are characteristic inflammatory biomarkers and hallmark of bacterial infection, respectively.

Quantification of CRP levels was performed at a private clinical laboratory (Martin Inc, Bayamón, PR) using an Architect c8000 Clinical Chemical Analyzer (Abbot, Illinois, US). PCT assay was performed using a Human Procalcitonin ELISA kit following manufacturer’s instructions (Abcam, UK, ab100630).

### Flow cytometry

To determine the effect of Fh15 on cells of innate immune system we used flow cytometry. A total of 150μl of anticoagulated peripheral blood was stained with surface antibodies cocktail for 30 minutes at 4°C in the dark. Two antibody cocktails were used for the analysis, which are listed in detail in the **Table 1S**. After cell staining, BD lysis buffer (BD Biosciences, USA) was used to lyse the red blood cells and fix the samples for 10 minutes in the dark. Cells were washed with FACS buffer three times. Samples were stored at 4°C in dark until the next morning for data acquisition using a MacQuant10 (Miltenyi Biotec, USA). Data was analyzed using FlowJo 10 (BD Biosciences, USA). Our gating strategy for monocytes based on their expression of CD14 and CD16 as described elsewhere (25) as follow: classical monocytes (CD14^++^, CD16^-^), non-classical monocytes (CD14^+^, CD16^++^) and intermediate monocytes (CD14^+^, CD16^+^). Neutrophils were gated based on their expression of HLA-DR, CD3^-^ and CD66 a/c/e^+^, dendritic cells (DCs) and plasmacytoid DCs based on their expression HLA-DR, CD11c^+^, CD3^-^, CD123^+^, and NK-cells based on their expression of CD3^-^, HLA-DR, CD8^+^ and NKG2a^+^.

### Sandwich ELISA for detecting circulating Fh15 in plasma

A sandwich ELISA was optimized to detect circulating Fh15 in plasma from animals that received an i.v. isotonic infusion containing Fh15. The assay uses the purified rabbit anti-Fh15 IgG and the anti-Fh15 IgG-HRP conjugate obtained as described above. Optimal concentration of the anti-Fh15 IgG as capturing antibody and the optimal dilution of the anti-Fh15-IgG-HRP conjugate were determined by ELISA checkerboard titration. The capturing antibody was assayed in duplicated using dilutions ranging among 0.25 to 20μg/ml in coating buffer (0.05M carbonate-bicarbonate pH 9.6). Disposable polystyrene 96-well plates (Costar, Corning, NY) were coated with 100μl/well of each capturing antibody dilution. After an overnight incubation at 4°C in a humid chamber the plate was washed three times with PBST and blocked with 5% skimmed milk-PBST (300μl/well) for 1 hour at 37°C in a humid chamber. After removing the blocking solution, undiluted plasma samples (100μl/well) were added to the plate and incubated by 1h 37°C after which plates was washed three times with PBST. Anti-Fh15 IgG-HRP at different dilutions ranging among 1:1,000 and 1:3,000 in PBST was added (100μl/well) and the incubation was prolonged for 30 minutes at 37 °C in humid chamber. After another washing step, the substrate solution (50ml 0.05M citrate-phosphate pH 5.0 + 20 mg *o*-phenylenediamine + 20μl H_2_O_2_) was added to each well (100μl/well) and the plate was incubated at room temperature for 15 minutes in the dark. The reaction was stopped by adding 50μl/well of 10% HCl and the optical density (OD) measured at 490nm using a SpectraMax M3 (Molecular Devices, LLC, USA). A standard curve was generated by adding Fh15 to a negative baseline plasma sample with different concentrations of Fh15 ranging from (1μg/ml to 61.03pg/ml). The OD at 490nm of spiked negative plasma sample was plotted against its known antigen protein concentration. The standard curve had an r^2^=0.99 using a 4 Parameter Logistic (PL) analysis. By using the standard curve, the ODs of plasma from animals used in the sepsis model were transformed into antigen concentrations (μg/ml).

### Plasma cytokines and chemokines level determination

Levels of plasma pro-inflammatory cytokines IL-6, IL-12p70, TNF-α, IFN-γ, and chemokines MCP-1, and IP10 were determined using a Non-Human Primate Customized Multiplex (R&D Systems, Minneapolis, USA) following manufacturer’s instruction by using a Magpix (Luminex, USA) and analyzed with the Bio-Plex Data Pro Software (BioRad, Hercules, CA).

### Statistical Analysis

All data was expressed as mean value ± S.D. for each determination. The statistical significance was determined using two-way ANOVA, Student t-test and non-parametric tests using GraphPad Prism software (version 8). For all tests, a *p* value < 0.05 was considered significant.

## Results

### Physiological parameters before and after sepsis induction

At baseline, BT, HR, and RR of all rhesus monkeys involved in the study were at normal values ranging between 33.4-37.2°C (median 35.9°C), 85-160 beats/min (median 114beats/min) and 13-25 breaths/min (median 21breaths/min), respectively. The MAP ranged between 33-78mmHg (median 51mmHg). These MAP measurements were lower than expected. According to recent literature oscillometric monitoring of BP are often underestimated (26, 27). At the conclusion of the procedure (8h) the physiological parameters were similar or slightly higher than the baseline for most of animals, which were euthanized without signs of septic shock

(**Table 1**). In contrast, for animal MA035 (group Fh15-Fh15-*E. coli*) MAP dropped from 41 to 21mmHg and the RR dropped from 20 to 5 breaths per minute (bpm). Despite this abrupt drop of RR, the animal did not die prematurely. Postmortem examination revealed no gross abnormalities with exception of splenomegaly, which was noticed in MA035, and lymphadenopathy that was observed in all animals from the *E. coli* control group.

**TABLE 1.** Main vital signs monitored in *Rhesus macaques* during 8 hours of experimental sepsis induced by intravenous infusion with live *E. coli*.

| Experimental Group | ID | Physiological Parameter | Baseline Value | Value at 8h of *E. coli* infusion |
| --- | --- | --- | --- | --- |
| *E. coli* | 6R1 | Body Temperature (°C) | 35.9 | 38.6 |
|  |  | Heart Rate (bpm) | 108 | 192 |
|  |  | Mean Arterial Pressure | 58 | 25 |
|  |  | Respiratory Rate (rpm) | 22 | 29 |
| *E. coli* | 0R5 | Body Temperature (°C) | 37.2 | 36.1 |
|  |  | Heart Rate (bpm) | 160 | 192 |
|  |  | Mean Arterial Pressure | 51 | 49 |
|  |  | Respiratory Rate (rpm) | 21 | 15 |
| *E. coli* | CB22* | Body Temperature (°C) | 36.3 | 37 |
|  |  | Heart Rate (bpm) | 124 | 172 |
|  |  | Mean Arterial Pressure | 31 | 71 |
|  |  | Respiratory Rate (rpm) | 16 | 24 |
| Fh15 | MA078 | Body Temperature (°C) | 35.7 | 36.4 |
|  |  | Heart Rate (bpm) | 111 | 130 |
|  |  | Mean Arterial Pressure | 50 | 55 |
|  |  | Respiratory Rate (rpm) | 21 | 21 |
| Fh15 | MA014 | Body Temperature (°C) | 36.8 | 35.9 |
|  |  | Heart Rate (bpm) | 148 | 130 |
|  |  | Mean Arterial Pressure | 78 | 68 |
|  |  | Respiratory Rate (rpm) | 14 | 24 |
| Fh15 | MA035 | Body Temperature (°C) | 37.2 | 37.2 |
|  |  | Heart Rate (bpm) | 140 | 142 |
|  |  | Mean Arterial Pressure | 53 | 70 |
|  |  | Respiratory Rate (rpm) | 25 | 30 |
| Fh15-Fh15-*E. coli* | MA078 | Body Temperature (°C) | 35.4 | 36.3 |
|  |  | Heart Rate (bpm) | 112 | 128 |
|  |  | Mean Arterial Pressure | 51 | 49 |
|  |  | Respiratory Rate (rpm) | 13 | 18 |
| Fh15-Fh15-*E. coli* | MA014 | Body Temperature (°C) | 33.4 | 34 |
|  |  | Heart Rate (bpm) | 120 | 164 |
|  |  | Mean Arterial Pressure | 33 | 20 |
|  |  | Respiratory Rate (rpm) | 23 | 20 |
| Fh15-Fh15-*E. coli* | MA035 | Body Temperature (°C) | 35.6 | 34.6 |
|  |  | Heart Rate (bpm) | 123 | 108 |
|  |  | Mean Arterial Pressure | 41 | 21 |
|  |  | Respiratory Rate (rpm) | 20 | 5 |
| Fh15-*E. coli* | 8Y4 | Body Temperature (°C) | 35.9 | 37.2 |
|  |  | Heart Rate (bpm) | 102 | 160 |
|  |  | Mean Arterial Pressure | 44 | 23 |
|  |  | Respiratory Rate (rpm) | 22 | 14 |
| Fh15-*E. coli* | 0Y2 | Body Temperature (°C) | 35.6 | 35.6 |
|  |  | Heart Rate (bpm) | 85 | 178 |
|  |  | Mean Arterial Pressure | 55 | 10 |
|  |  | Respiratory Rate (rpm) | 22 | 22 |
| Fh15-*E. coli* | IZ8 | Body Temperature (°C) | 36.1 | 34.8 |
|  |  | Heart Rate (bpm) | 114 | 156 |
|  |  | Mean Arterial Pressure | 36 | 24 |
|  |  | Respiratory Rate (rpm) | 21 | 21 |

\**Rhesus* from the study published in *Animal Model Exp. Med.* 2019; 2:326-333

### Developing of antibodies to Fh15 and dynamic of Fh15-antigenemia

A double antibody sandwich ELISA optimized as described above was used to measure levels of circulating Fh15 in plasma of Fh15-infused rhesus macaques. Antibodies against Fh15 were measured by a previously optimized indirect ELISA (22). As expected, any of naïve rhesus involved in the study had antibody against Fh15 or circulating Fh15 in the plasma samples collected at baseline, which confirmed that none of animals had been previously exposed to this antigen or were infected with parasites that potentially could induce cross-reactive antibodies to Fh15.

When the plasma samples collected from animals that only received Fh15 (MA014, MA035 and MA078) were tested for antigenemia we found maximal Fh15 concentrations (3.48±3.31μg/ml) at 30 min following the infusion, which at 2h declined to an average of 0.135±0.048μg/ml and undetectable at subsequent time points. In the rhesus that received the Fh15 + *E. coli*-infusion (Fh15-*E. coli*, 8Y4, 0Y2, IZ8) the Fh15 average concentration at 30 min was 6.12-fold lower than in the Fh15-group (mean 0.568±0.11μg/ml). At 2h the antigenemia declined in this group to an average of 0.179±0.067μg/ml and was undetectable thereafter. Interestingly, the group termed Fh15-Fh15-*E. coli* (comprised by the rhesus MA014, M035 and MA078 that had received the Fh15-infusion 3 months before) showed the most intense and prolonged antigenemia. The maximal Fh15 average concentration at 30 min was 7.53±3.44μg/ml, which is 2.16-fold higher than those observed in the Fh15-group and 13.25-fold higher than the observed in the Fh15-*E. coli* group. At 2h the antigenemia dropped to an average of 3.599±0.95μg/ml, at 4h to 1.685±0.612μg/ml and at 6h still was detectable with concentrations of 0.94±0.21μg/ml. After 8h, circulating Fh15 was no longer detected (**Figure-1A**). However, the plasma sample collected from these rhesus at baseline of the second Fh15-infusion tested positive for antibodies with titters against Fh15 among ∼1:200 to 1:400.

**Figure-1.**
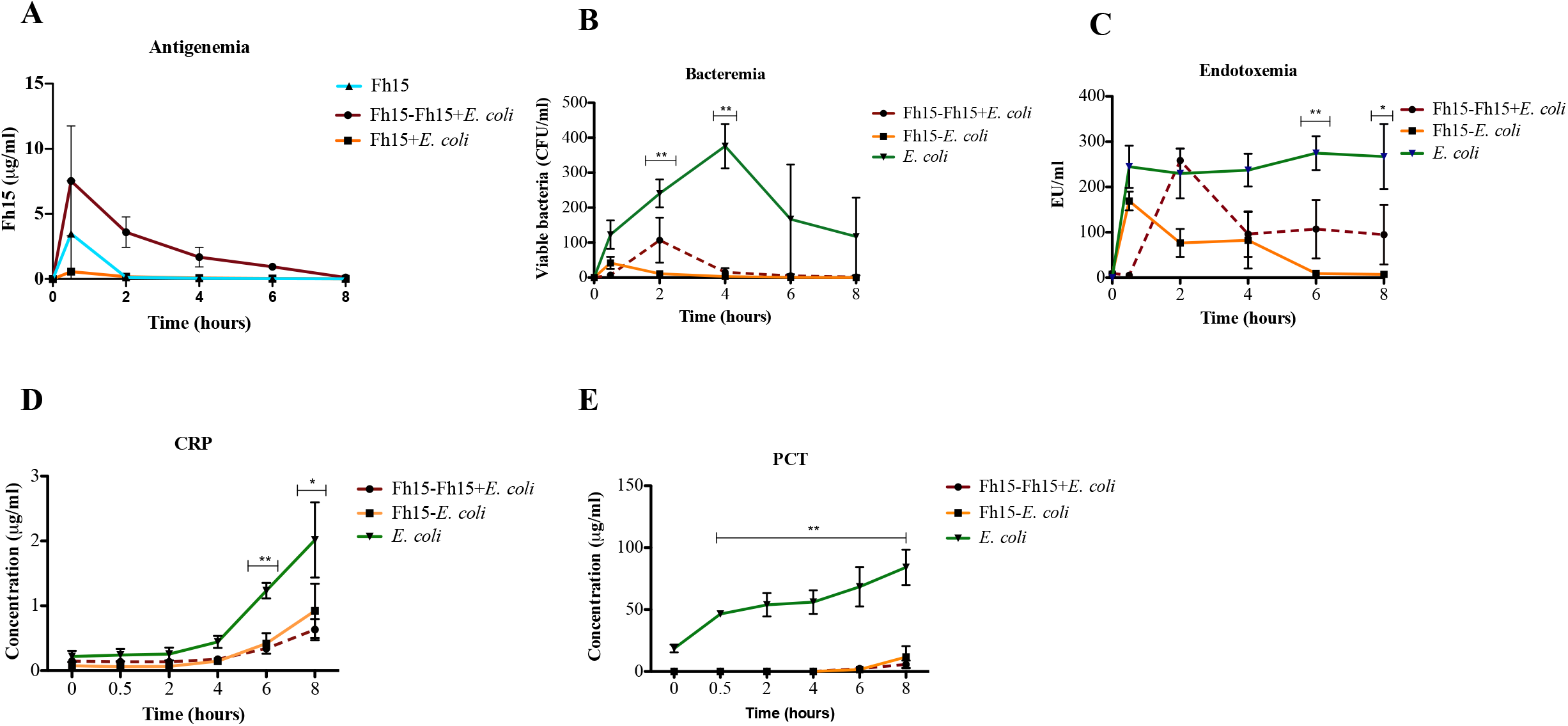
Fh15 decreases bacteremia, endotoxemia, C-reactive protein and Procalcitonin in rhesus during an acute lethal endotoxemia. The graphs represents the dynamic of levels of **(A)** Fh15 antigenemia, **(B)** viable bacteria (\*\**p*=0.0046 & *p*=0.0042), **(C)** plasma endotoxin (LPS) (\*\**p* =0.0021 & *p*=0.0224), (**D**) C-reactive protein (\*\**p*= 0.001) and **(E)** procalcitonin (\*\**p* =0.001) over time in groups (n=3) of male rhesus that only received the bacterial infusion (*E. coli*), Fh15 plus *E. coli*-infusion (Fh15-*E. coli*) or two doses of Fh15 three months apart + *E. coli* (Fh15-Fh15-*E. coli*). Statistical significance between control group and experimental groups was determining by ANOVA or Student t-test using GraphPad Prism-8. For all tests, a *p* value < 0.05 was considered significant.

### Fh15 suppress the bacteremia, endotoxemia and the production of acute-phase proteins

In agreement with our previous observations using the rhesus macaque model of sepsis, (20) rhesus that only received the bacterial infusion (*E. coli*: 6R1, 0R5, CB22) had high levels of bacteremia, which were notably high from 30 min following the bacterial infusion administration (123±57.3 CFU/ml). Overall, the bacteremia in this group increased progressively until reaching a peak at 4h (376±89CFU/ml). Then declined and remained at detectable levels during the full-time course. One animal of this group (BP22) had high and persistent levels of bacteremia at every time point (average 385±62.24CFU/ml) whereas in the other two animals (6R1 and OR5) the average number of viable bacteria declined dramatically to 10±2CFU/ml and 4CFU/ml at 6h and 8h, respectively (**Figure 1B**). The rapid decline of bacteremia in two animals indicate that bacteria were unable to colonize and replicate inside these animals and were lysed by the complement system as it has been reported in experimental conditions where high bacteria inoculum are used to induce sepsis (28). Since all animals received the same bacterial dose adjusted to the body weight, the detection of bacteremia in one animal at every time point suggests failure in the efficiency of the complement system to lyse bacteria.

As expected, the levels of plasma LPS in all animals of the *E. coli*-group increased abruptly at 30 min (244.81±65.76EU/ml) and remained at very high levels throughout the experiment (252.3±76.8EU/ml) (**Figure 1C**). The increase of plasma endotoxin is a logical consequence of the bacteria disruption and consequent releasing of LPS to the bloodstream, an observation that also was seem in our previous study developing the rhesus model of endotoxemia (20). The C-reactive protein (CRP) (**Figure 1D**) an acute phase protein that typically increases in plasma during inflammatory process (29) was detectable at 6h and reached maximal values at 8h following the *E. coli*-infusion (2.016±0.81μg/ml). Procalcitonin (PCT) (**Figure 1E**) a marker of bacterial infection and a predictor of sepsis in humans (30) was detected from 30 min and increased sequentially until reach maximal levels at 8h following *E. coli*-infusion (84.12±20.27ng/ml).

The animals that only received the Fh15-infusion did not developed bacteremia, endotoxemia, CRP or PCT at any time throughout the study (data not shown). This is clear evidence that the Fh15-infusion was prepared pyrogenic-free and administered to animals following good laboratory practices (GLP). An important finding of the present study is that the number of blood viable bacteria in the Fh15-*E. coli* group was drastically lowered at every time point compared to the *E. coli*-group (**Figure 1B**). These animals had 2.93-fold lower viable bacteria (average 42±24.3CFU/ml), 21.8-fold lower (average 11±6.37CFU/ml) and 125-fold lower (average 3±2.1CFU/ml) than in the *E. coli*-control at 30 min, 2h (*p*=0.0046) and 4h (*p*=0.0042), respectively, with no viable bacteria detected in the subsequent time points.

When we analyzed the bacteremia in rhesus from the Fh15-Fh15-*E. coli* group a similar pattern was observed although with some differences. In two animals (M014 and M035) the bacteremia reached maximal values at 2h (47±35CFU/ml) declining quickly at 4h (4CFU/ml) and making undetectable thereafter. Whereas in the rhesus MA078 the number of viable bacteria detected at 2h was high (226CFU/ml), declined to 38CFU/ml by 4h and to 16CFU/ml and 6CFU/ml at 6h and 8h, respectively. Statistical differences were found between the bacteremia in the Fh15-Fh15-*E. coli* group compared to the *E. coli*-group at 30 min (*p*=0.0488) and 4h (*p*=0.0055). However, no statistical differences were found between the levels of bacteremia in the Fh15-*E. coli* and Fh15-Fh15-*E. coli* groups at any of time points studied. These results demonstrate that the treatment with Fh15 applied 30min prior to exposure to a lethal live *E. coli* infusion can efficiently prevent the bacterial replication in blood.

In agreement with the decreasing of bacteremia, the LPS levels in plasma were also notably lower in both experimental groups compared to the *E. coli* control group. In the group Fh15-*E. coli* group average concentration of LPS at 30 min was 169.16±29.27EU/ml, which represents a significant lowering of 1.44-fold less LPS than in the *E. coli* control group (p=0.0067). The concentrations of LPS in this experimental group lowered subsequently reaching the lowest levels at 8h (7.3±2.5EU/ml), which represented a significant diminution (p=0.0042) of 28.24-fold compared to the *E. co*li-group (**Figure 1C**). In the group Fh15-Fh15-*E. coli* the levels of LPS at 30 min were very low but at 2h had reached a peak (average 258.33±38.11EU/ml). Then in two animals (M014 and M035) the LPS concentrations reduced sequentially until reach lowest levels of 5.5±0.45 EU/ml at 8h, which represented a significant lowering of 48.14-fold lower concentration than in the *E. coli* group (*p*=0.0224). However, in the rhesus MA078 the LPS concentrations remained consistently high throughout the experiment with an average concentration of 218.6±17.63EU/ml. Statistical differences between both experimental groups were found at 30 min (p=0.0014) and 2h (*p*=0.0114) and not in the subsequent time points.

Importantly, all animals from experimental groups (Fh15-*E. coli* and Fh15-Fh15-*E. coli*) had significantly lower levels of CRP and PCT than the *E. coli* control group (*p*=0.001) (**Figure 1D & E)**. In humans undergoing sepsis or septic shock, CRP is known to activate the complement system and increase antigen presentation. Also, it has been correlated with an increased organ dysfunction, longer hospital Intensive Care Unit (ICU) stay and mortality (31). PCT in the ICU is used as a biomarker for survival, with lower levels resulting in lower sequential organ failure assessment (SOFA) scores and better prognosis in septic patients (32).

### Fh15 suppress the production of pro-inflammatory cytokines in plasma of septic rhesus macaques

Having demonstrated that the administration of an Fh15-infusion prior to a live bacterial infusion is able to suppress bacteremia, endotoxemia as well as levels of CRP and PCT induced by *E. coli*, we proceeded to investigate whether Fh15 could also suppress several pro-inflammatory cytokine/chemokines that are signatures of inflammation during sepsis. As expected, all animals from the *E. coli* group developed a strong pro-inflammatory cytokine storm evidenced by the high levels of IFNγ, IL-6, TNFα, IL-12, IP-10, and MCP-1 throughout the entire time-course. This is consistent with the high levels of LPS, CRP and PCT detected in plasma of these animals, which is indicative of an ongoing endotoxemia. However, all animals from groups Fh15-*E. coli* and Fh15-Fh15-*E. coli* had lower concentrations of IL-6, TNFα, IL12, IFNγ cytokines and lower IP10 and MCP-1 chemokines than animals from the *E. coli* control group at every time point studied (**Figure 2**). Unfortunately, due to the large immunological variability in levels of cytokines/chemokines among outbred individuals from the same group and the low number of animals within each group, no statistical differences were found. To better appreciate the reduction that produced Fh15 in the levels of these pro-inflammatory cytokines/chemokines we analyzed the ratio in the average concentration of each cytokine/chemokine in the *E. coli* control group compared to the Fh15-*E. coli* and Fh15-Fh15-*E coli* groups (**Figure-3S**). This analysis revealed that animals from the *E. coli*-control group had 1.3 to 24-fold more IFNγ than those from the Fh15-*E. coli* group at 2h and 8h, respectively. Similarly, the *E. coli* control group had 1.5 to 7.03-fold more IFNγ than the animals of Fh15-Fh15-*E. coli* group at the same time-points. IFNγ is a cytokine classically produced by NK-cells and T lymphocytes, which facilitates systemic inflammation during endotoxin-induced shock (33). IL-12, a cytokine naturally produced by DCs, macrophages and neutrophils cells in response to antigenic stimulation (34-38) was found 75.8-fold to 386.2-fold more increased in the *E. coli-*control group than in the Fh15-*E. coli* group and was found 23.3-fold to 224-fold more increased in the *E. coli*-control group than in the Fh15-Fh15-*E. coli* group. TNFα an inflammatory cytokine produced by macrophages/monocytes during acute inflammation (39), which normally has a peak of secretion about 2h following the *E. coli* insult (20) with a further decline later was found 23.34-fold and 6.56-fold more increased in the *E. coli*-control group than Fh15-*E. coli* group and Fh15-Fh15-*E. coli* group, respectively. After declining, the *E. coli*-control group maintained a ratio of >1.4-fold more TNFα than the experimental groups throughout the entire time-course. IL-6, a cytokine produced by a variety of cells including macrophages and monocytes during inflammatory process (40) was also found 5.06-fold more increased at 2h in the *E. coli* control group than in the Fh15-*E. coli* and 12.6-fold more increased than in the Fh15-Fh15-*E. coli* group. The *E. coli* control group maintained a ratio >1.5 fold more IL-6 than the experimental groups throughout the other time points. The same observations were noticed for chemokines IP-10 and MCP-1, which were found 5 to 2.8-fold and 2.08 to 3.63-fold more increase in the *E. coli* control group than experimental groups, respectively.

**Figure 2.**
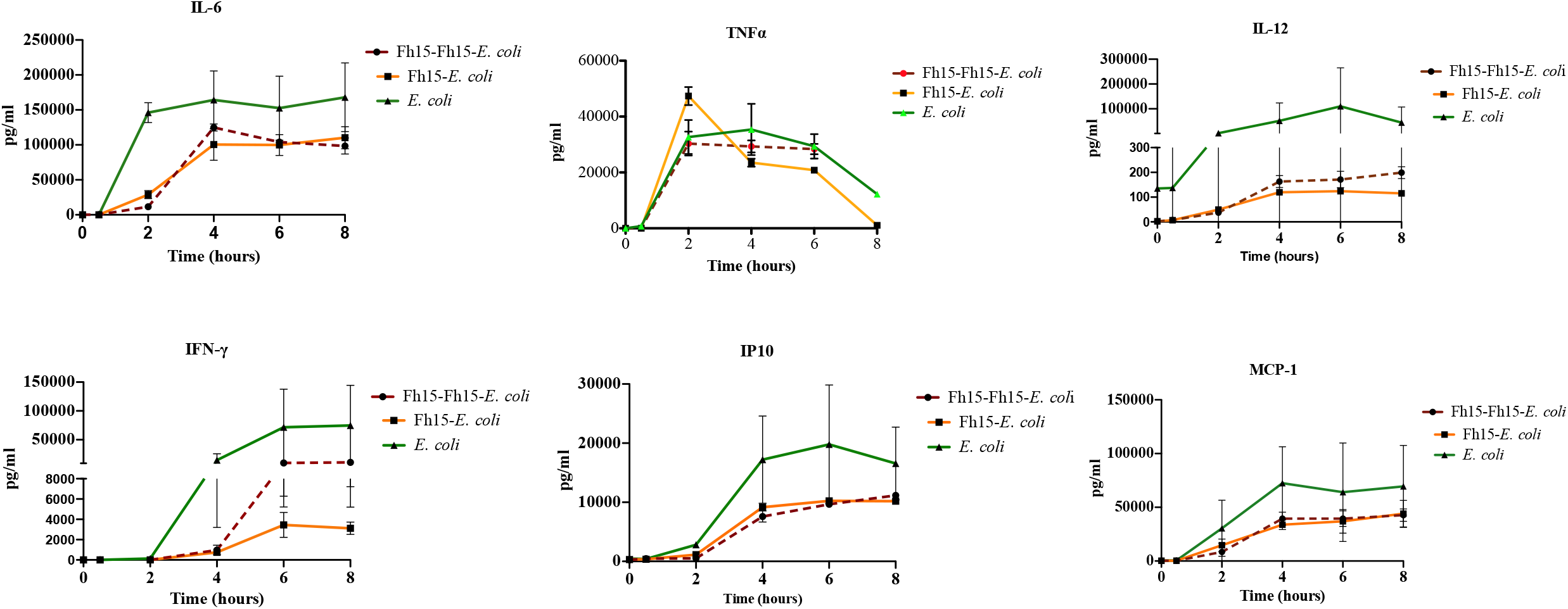
Fh15 decreases the production of pro-inflammatory cytokines and chemokines in rhesus undergoing sepsis. Plasma samples were collected from rhesus macaques at different time points: 0 (baseline), 30 min, 2h, 4h, 6h and 8h. Animals were allotted into three experimental groups (n=3 each) termed ***E. coli***: control group that only received the i.v. infusion with live *E. coli* (10^10^ CFU/kg body wt.), **Fh15-*E. coli***: received an i.v. infusion containing 12mg Fh15 + *E. coli*-infusion and **Fh15-Fh15-*E. coli***: received an i.v. infusion with 12mg Fh15 three months before and a second Fh15-infusion + *E. coli*-infusion. Levels of IFN-γ, IL-6, IL-12, TNFα, MCP-1 and IP10 were measured using Luminex technology.

### Effect of bacterial infusion on the innate immune cells population and the counter effect caused by Fh15

When the blood innate immune cell populations in the group that only received the *E. coli* infusion was examined it was noticed that most of the cell populations had significantly decreased by 30 min following the infusion (*p*<0.0001) and remained at very low levels throughout the entire full-time course (**Figure 3**). Specifically, peripheral blood monocytes dropped >14.8-fold compared to baseline. Classical monocytes, which comprise about 80-95% of circulating monocytes and are highly phagocytic (41, 42) had dropped >1,000-fold. Non-classical monocytes that comprise about the 2-11% of circulating monocytes (25) and have pro-inflammatory behavior had dropped >100-fold. Intermediate monocytes, which comprise about 2-8% of circulating monocytes and have among their functions the production of reactive oxygen species (ROS), antigen presentation, stimulation and proliferation of T-cells and angiogenesis (25) had dropped >40-fold. Dendritic cells (DCs) and plasmacytoid DCs (pDCs), which play pivotal roles in the initiation of innate and adaptive immune response to pathogens (43) dropped >50-fold and > 6-fold, respectively. Natural killer (NK)-cells, which make up 5-15% of human peripheral blood (44) and play protective roles against both infectious pathogens and cancer (45) dropped >12-fold.

**Figure-3.**
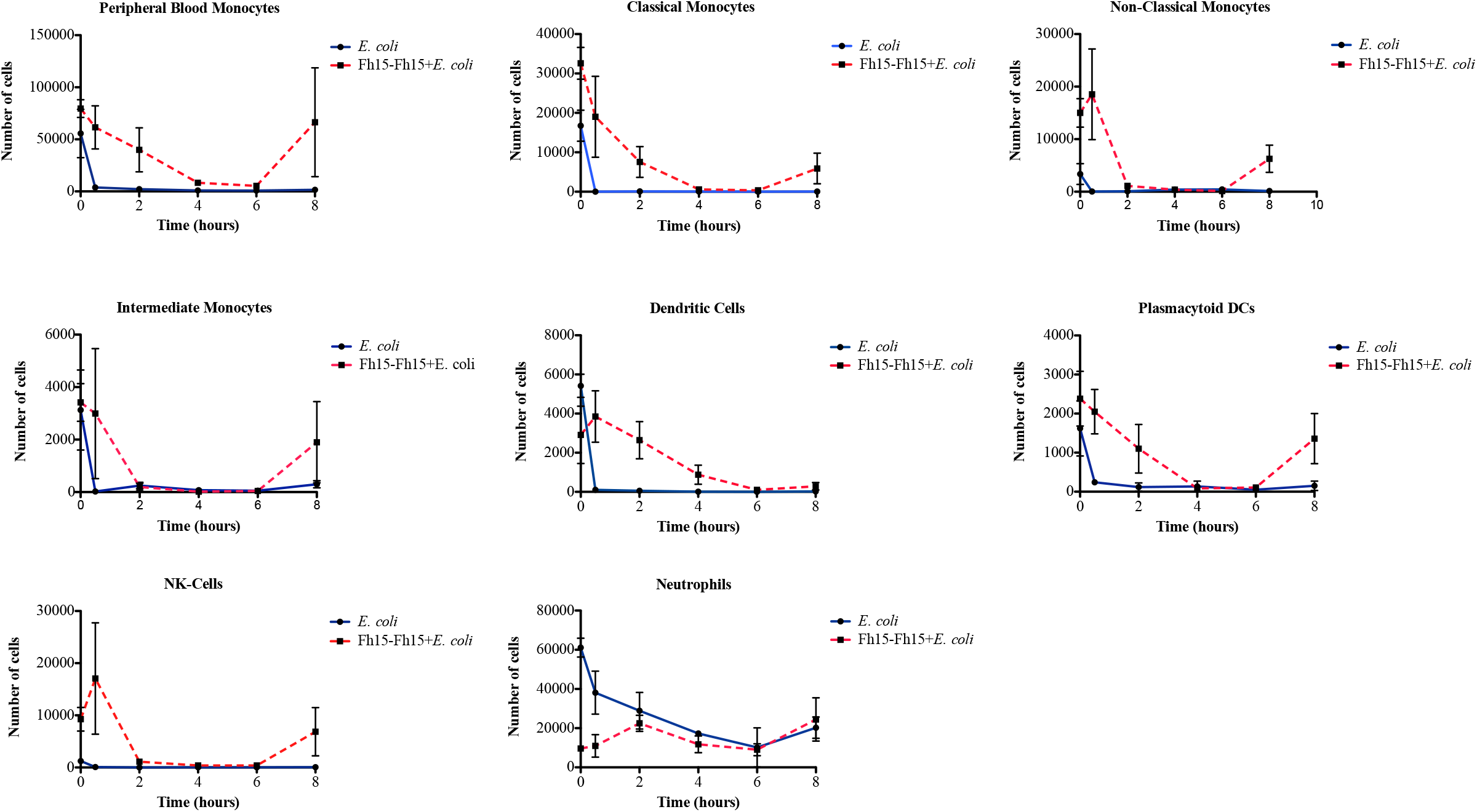
Effect of live *E. coli* compared to two Fh15-doses administered three months apart, prior to *E. coli*-infusion on the number of innate immune cells in the bloodstream. Graphs represent the number of peripheral blood monocytes, classical monocytes, non-classical monocytes, intermediate monocytes, dendritic cells, plasmacytoid DCs, natural killer cells and neutrophils over time in two experimental groups: *E. coli*: rhesus (n=3) that received an i.v. isotonic infusion with live *E. coli* (10^10^ CFU/kg body wt.). Fh15-Fh15-*E. coli*: rhesus (n=3) that received an i.v. infusion with 12mg Fh15 and 3 months later they received a second Fh15-infusion (12mg) followed by the *E. coli*-infusion. Whole blood samples were stained with a cocktail of antibodies for labeling surface markers specific for peripheral monocytes (CD14^+,^ CD16^+^), myeloid and plasmacytoid dendritic cells (CD11c^+^, CD123^+^), natural killer cells (HLA-DR^+^, CD3^-^, CD69^+^) and neutrophils (CD66a/c/e^+^). Cells were analyzed by gating using a MacQuant10. Data was analyzed using FlowJo. Data represent average number of cells ± SD of three independent biological samples, each in duplicate at every time point studied.

Interestingly, in the group that only received the Fh15 infusion most these cell populations increased progressively until reaching maximal levels between 4 to 6 hours. At these time points the amount of classical monocytes population was 35-fold higher than baseline, there was 1.44-fold more non-classical monocytes, and 1.18-fold more intermediate monocytes than in the baseline. The DCs and NK-cells populations also were found 2.13-fold and 1.6-fold more abundant than baseline, respectively. Although the number of cells decreased slightly after peak all cell populations remained at high levels until the end of the study. The pDCs and neutrophils remained at levels relatively like baseline although the trend of pDCs was to decline sequentially. Importantly, a dynamic almost identical to those described for Fh15 was observed within animals from the Fh15-*E. coli* group (**Figure 4**). The Fh15-Fh15-*E. coli* group had a different dynamic of innate immune cells in which the amount of all monocyte populations, DCs, and NK-cells declined sequentially until reach the lowest levels amount between 4 to 6 hours and then increased notably at the end of the procedure **(Figure 3**).

**Figure-4.**
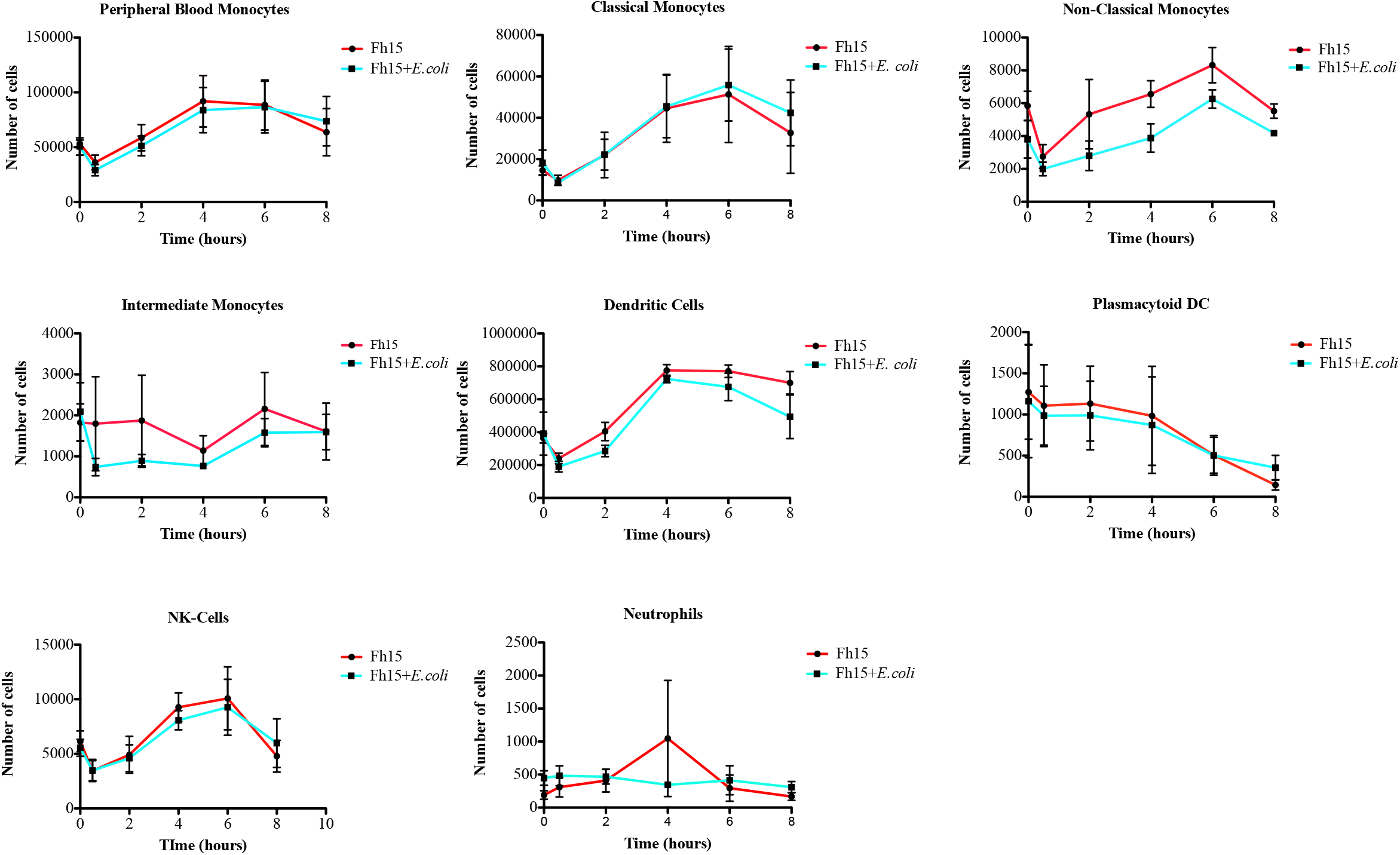
Fh15 treatment increases the innate immune cells in blood stream. Graph represents the number of peripheral blood monocytes, classical monocytes, non-classical monocytes, intermediate monocytes, dendritic cells, plasmacytoid DCs, natural killer cells and neutrophil over time in two experimental groups. Fh15: a group of animals (n=3) that only received the Fh15 infusion (12mg) and Fh15-*E. coli*: a group of animals (n=3) that received the Fh15-infusion followed by the *E. coli* infusion. Whole blood samples were stained with a cocktail of antibodies for labeling surface markers specific for peripheral monocytes (CD14^+,^ CD16^+^), myeloid and plasmacytoid dendritic cells (CD11c^+^, CD123^+^), natural killer cells (HLA-DR^+^, CD3, CD69^+^) and neutrophils (CD66a/c/e^+^). Cells were analyzed by gating using a MacQuant10. Data was analyzed using FlowJo. Data represent average number of cells ± SD of three independent biological samples, each in duplicate, at every time point studied.

## Discussion

Our previous *in vitro* studies using THP1-Blue CD14 cells and mouse bone marrow-derived macrophages demonstrated that Fh15 can suppress the NF-κB activation induced by different Gram-negative and Gram-positive bacteria extracts and suppress a number of the LPS-induced pro-inflammatory cytokines, respectively (22). Moreover, *in vivo* studies using a mouse model of sepsis demonstrated that Fh15 could be useful as prophylactic and therapeutic drug to prevent the dissemination of the cytokine storm in the animals that received Fh15 (18, 23). Based on these antecedents and as proof of concept, the current study aimed to ascertain whether Fh15 could be useful limiting the lethal inflammatory responses and bacterial dissemination caused by an i.v. infusion with live *E. coli* in rhesus macaques.

Fh15 is an immunogenic molecule capable to induce strong antibody response when is injected subcutaneous in rabbits or mice emulsified in adjuvant (16, 46). Hence, rabbits and mice infected with *F. hepatica* also develop antibodies against Fh15 (16, 46). Therefore, it was not surprising that all animals that received the Fh15-infusion have detectable antibody against Fh15 three months later. However, what was surprising was the observation that these same animals not only had antibodies, but also higher levels of antigenemia. The detection of antibodies against Fh15, concurrently with circulating Fh15, suggests that the antibodies elicited against Fh15 had low affinity and did not attach strongly enough to the antigen; otherwise, the sandwich ELISA would not have detected circulating Fh15. The production of antibodies of low titter and low affinity often occurs when the exposure to the antigen has been discontinued and fail the selection of clones that produce antibody of high affinity (47).

The rationale to include in the present study an experimental group that has antibodies specific to Fh15 was precisely to determine whether these antibodies may block the ability of Fh15 to exert its anti-inflammatory function. In that regard, it was interesting to observe that animals with detectable anti-Fh15 antibodies (Fh15-Fh15-*E. coli* group) had similar ability to those that were naïve to Fh15 (Fh15-*E. coli*), to prevent bacterial replication, reduce the concentration of plasma LPS and suppress the production of CRP and PCT (**Figure 1**), suggesting that at least these functions were not impaired by the presence of antibodies.

Although the mechanism that uses Fh15 to prevent the bacterial replication remains unknown we have speculated that Fh15 could be inducing immune cells to release of extracellular DNA traps (ETs) to trap and kill bacteria. This process termed Etosis is a distinct process of cell death and has been reported occur in several immune cells including neutrophils (48), eosinophils (49), mast cells (50) and monocytes/macrophages (51) from humans or mice. This hypothesis rose from a preliminary experiment in which blood monocytes isolated from naïve rhesus macaques were differentiated to macrophages *in vitro* (52) and then treated for 16h with LPS (100ng/ml), Fh15 (20μg/ml), LPS + Fh15 or DNAseI (1mM). It was observed that cells incubated with Fh15 and/or LPS detached and showed a necrotic appearance whereas cells incubated with DNAseI showed a healthy appearance. Thus, the population of phagocytic cells in the bloodstream, which were notably increased in the presence of Fh15 (**Figure 4**) could be releasing extracellular DNA traps and antimicrobicidal proteins to enhance bacteria killing as has been already reported occurs in the presence of *E. coli* (53) or fungi such as *Candida albicans* (54). Etosis has been also reported in humans and mouse immune cells either *in vivo* or *in vitro* in response to protozoa and helminth parasites infections including *Toxoplasma gondii (55), Leishmania amazonensis* (56) and *Strongyloides stercoralis* (57). However, this pathogen killing mechanism has not been reported in rhesus macaques so far and there are no reports demonstrating that it can be induced by injections with parasite-derived antigens. Further experiments should be done to confirm that Etosis is the mechanism of action by which Fh15 reduces bacteremia and endotoxemia in this model.

Another interesting observation of the present study was the abrupt and sustained decrease in the blood innate immune cells during the 8 hours following to the *E. coli*-infusion (**Figure 3**). Blood monocytes (including classical, non-classical and intermediate monocytes), NK cells and DCs are key cells during early phase of endotoxemia because they are not only responsible for maintaining vascular homeostasis, but also are highly responsible for patrolling the bloodstream to recognize and phagocyte invading pathogens, resulting in the secretion of pro-inflammatory cytokines, such as IL-1β, TNF-α, IL-6 and IL-8 (58-60). The ‘disappearance’ of immune cells from the blood stream in rhesus from the *E. coli*-group could be a logical consequence after an inflammatory insult since immune cells migrate to secondary lymphoid organs to maturate, differentiate, and present the antigen to the naïve T-cells (61). This observation is consistent with a similar phenomenon fully documented that occur in mice exposed to lethal intraperitoneal doses of LPS in which large peritoneal macrophages (LPMs), the macrophage population more abundant at steady state, “disappear” from the peritoneal cavity at the beginning of the LPS-insult (18, 62). In that regard, we have demonstrated that Fh15 shifts the LPMs dynamic by not only preventing the disappearance of LPMs but also augmenting this cell population in the peritoneal cavity (18). The observation that Fh15 alone or in the presence of *E. coli* can promote the persistence of innate immune cells in the bloodstream during the early phase of the endotoxemia leads us to suggest that a primary modulatory mechanism of Fh15 would be based on generating an immunological environment compatible with the homeostasis or a prolonged steady state even in the presence of inflammatory stimuli. However, based on the results showed in **Figure 3**, the ability for Fh15 to increase the innate immune cell population seems to be partially abrogated in the rhesus macaques exposed to Fh15 three months before and that had antibodies against Fh15 (Fh15-Fh15-*E. coli*). However, it should be noticed that the reduction of cell populations in this experimental group was sequential and not abrupt like occurred in animals that only received *E. coli* and at the end of the experimental window all cell populations had again increased. Therefore, considering that in these same animals the presence of antibodies does not seem to affect the decrease of bacteremia, endotoxemia, CRP or PCT or the ability for Fh15 to reduce the pro-inflammatory cytokine/chemokines **(Figure 2)**, it should not be ruled out that other immunologic factors, unknown so far, could have influenced this result.

The observation that Fh15 alone does not induce IL-12, IL-6, IFNγ, TNFα, IP-10 or MCP-1 is consistent with our previous studies in which native or recombinant *F. hepatica* FABPs are unable to elicit inflammatory responses (18, 22, 23). Importantly, despite the large variability among the cytokine/chemokine concentrations for each rhesus in the experimental groups, it was possible to observe a notable lowering in the concentrations of IL-12, IL-6, IFNγ, TNFα, IP-10 or MCP-1 in plasma of animals that received the Fh15-infusion compared to those that only received the *E. coli-*infusion. The lack of statistical differences in these determinations could be due to several factors affecting simultaneously. Due to the immunological variability typical in outbreeds subjects, the low number of animals per group, and the lack of an appropriate adjustment in the amount of Fh15 administered to rhesus according to their body weight made it difficult to determine the real behavior of data and remove outlier subjects. It should be highlighted that the dose of Fh15 administered to rhesus was arbitrarily selected based on the amount of purified Fh15 availability rather than a rigorous dose-dependent experiment to select the best Fh15 concentration that maximizes the results. If we consider that the groups of rhesus in our experiment was weighing 6.4 to 9.66kg and all received the same Fh15 dose (12mg), it meant that some animals received 1.87 mg/kg body wt. of Fh15 whereas others received a dose equivalent to 1.24mg/kg/body wt. Overall, the amount of Fh15 administered to rhesus in the present study was lower than those applied intraperitoneal to BALB/c mice (50μg per mouse) (18), which assuming that all mice weighting ∼20g that would be equivalent to 2,500μg/kg body wt. Moreover, as mentioned above, the observation that the group that had anti-Fh15 antibodies (Fh15-Fh15-*E. coli*) reduced the concentrations of plasma IL-12, IL-6, IFNγ, TNFα, IP-10 or MCP-1 in a similar magnitude to those observed in the Fh15-*E. coli* that does not have anti-Fh15 antibodies, indicate that the presence of anti-Fh15 antibodies do not impair the capacity of Fh15 to suppress the cytokine storm. This is consistent with a previous experiment performed with THP1-Blue CD14 cells in which it was demonstrated that the presence of anti-Fh12 antibodies do not abrogate the capacity of Fh15 to suppress the LPS-induced TLR4-stimulation (22). The lowering of pro-inflammatory cytokines induced by Fh15 is also in agreement with other reports in which *F. hepatica* infection has shown to suppress Th1 responses in concurrent bacterial infections (11, 63). These findings prelude a promising therapeutic outcome in rhesus pre-clinical sepsis model and support the therapeutic potential of Fh15 as an anti-inflammatory agent.

## CONCLUSIONS

Three novel issues stand out from the results being reported in this communication, that a small recombinant molecule belonging to the *F. hepatica* fatty acid binding protein (Fh15) administered intravenously in rhesus macaques prior to an infusion with lethal amounts of live *E. coli* a) significantly reduces bacteremia, suppresses the levels of LPS in plasma and blocks induction of CRP and PCT, which are key signature of inflammation and bacterial infection, respectively, b) notably reduced the production of pro-inflammatory cytokines and chemokines, and c) prevented the immune cell ‘disappearance’ from blood stream. Instead, Fh15, promoted the increase of innate immune cell populations in blood, a phenomenon that seems to be like those observed for LPMs in peritoneum of septic mice. This could suggest a role for Fh15 in promoting a prolonged steady state or homeostasis in rhesus even in the presence of inflammatory stimuli.

The small number of samples and the lack of an adequate adjustment in the Fh15 dose based on the body weight, which could have contributed to the variability in some results, limit the present study. However, despite these limitations, the results of this communication showing that a single prophylactic dose of Fh15 can suppress the bacteremia, the levels of endotoxemia as well as the levels of acute-phase proteins and a large panel of pro-inflammatory cytokines/chemokines during the early phase of sepsis is highly promising and could be translated to a better prognosis in this rhesus pre-clinical sepsis model. More pre-clinical studies are in progress using the rhesus model to determine if as hypothesize, Fh15 promotes the bacterial killing by Etosis, evaluate the effectiveness of Fh15 lowering inflammatory markers when is injected therapeutically after a live *E. coli* infusion or after an ongoing sepsis and importantly, defining the optimal Fh15 dose that maximize the results.

## ACKNOWLEDGMENTS

This research was supported by the National Institute of Allergy and Infectious Diseases Grant/Award Number: 1SC1AI155439-01, Office of Research Infrastructure Program of NIH (ORIP-NIH) 2016-2021, Grant/Award Number: P40OD12271; National Institute on Minorities Health and Health Disparities, Gran/Award Number: 5R25GM061151, G12MD007600 and R25GM061838. The content is solely the responsibility of the authors and does not necessarily represent the official views of the National Institutes of Health. Authors wish to thank Dr. Carlos A. Sariol (Director of Comparative Medicine) and Dr. Melween Martinez (Director of the Caribbean Primate Research center of Puerto Rico) for providing the animals and all the logistic to support to accomplish this study.

## AUTHOR CONTRIBUTIONS STATEMENT

The authors declare no conflict of interest.

**JJRF:** Participated in the experimental design, performed all experiments, the statistical analysis, prepared figures and revised the manuscript.

**AAR:** Collaborated in the samples processing and cytokine/chemokine determinations and revised the manuscript.

**NMR:** Collaborated in the flow cytometry experiments and data analysis, and revised the manuscript

**AB & NS:** Handled the animals, administered the i.v. infusions and collected the blood samples **SDE**: Provided advisory in the flow cytometry gating strategy and data analysis, and revised the manuscript

**LBM:** Provided BioPlex technology, collaborated in the cytokine analysis and revised the manuscript

**AME:** As PI, provided the funding and laboratory space where the experiments were performed. Participated in the experimental design, participated in the data analysis, prepared the figures, and wrote the manuscript.

## LEGENDS OF FIGURES

**Figure 1S. Purity of Fh15**. Recombinant Fh15 was successfully expressed within *Bacillus subtillis* as fusion protein with 6His-Tag at the amino terminal. SDS-PAGE and Western Blot were used to assess the purity of the protein. Lane-1 represents a BSA (2μg) as protein control, lane-2 represents Fh15 (2μg) under reducing conditions and lane-3 represents Fh15 (2μg) under reduced conditions incubated with a mouse antibody against the histidine tag.

**Figure 2S. Schematic representation of the experimental design**. Diagram represents the experimental design performed with rhesus macaques. Experimental groups (n=3) comprised naïve male rhesus average weighting ∼ 8.46kg. Animals were allotted into four groups. The control group termed *E. coli*, only received an i.v. isotonic infusion (50-ml) containing a lethal dose of live *E. coli* (10^10^ CFU/kg body wt.). This infusion was administered by 2h at a flow rate ∼0.416ml/min. Another group termed Fh15 only received the isotonic infusion (5-ml) containing 12mg Fh15. This infusion was administered during 20 min at a flow rate ∼0.25ml/min. The group termed Fh15-*E. coli* received first the Fh15-infusion followed by the *E. coli*-infusion. The group termed Fh15-Fh15-*E. coli* was comprised by the same animals from Fh15 group, which 3 months after receiving the first Fh15 infusion were returned to the experiment to receive a second Fh15-infusion + *E. coli* infusion. Blood samples were collected at baseline (0 min), 30 min, 2h, 4h, 6h and 8h. All animals that received the *E. coli* infusion were euthanized at 8h. Blood samples collected were used for determining levels of bacteremia and then staining with two antibody cocktails specific for cell markers of innate immune cell populations. These samples were analyzed by flow cytometry. Plasma recovered from the blood samples were used for measuring levels of LPS, antigenemia, CRP, PCT and cytokines/chemokines. Figure was created in BioRender.com

**Figure 3S. Ratio of cytokine/chemokine for each experimental group**. This graph represents the ratio between the average concentration (pg/ml) for each cytokine or chemokine in the *E. coli* control group compared to the average concentration for each cytokine/chemokine in the Fh15-*E. coli* or Fh15-Fh15-*E. coli* groups during the full-time course.

